# Differential contribution of direct and indirect pathways from dorsolateral and dorsomedial striatum to motor symptoms in Huntington’s disease mice

**DOI:** 10.1101/2024.06.12.597534

**Authors:** Sara Conde-Berriozabal, Laia Sitjà-Roqueta, Esther García-García, Lia García-Gilabert, Anna Sancho-Balsells, Sara Fernández-García, Ened Rodriguez-Urgellés, Albert Giralt, Javier López-Gil, Emma Muñoz-Moreno, Anna Castañé, Guadalupe Soria, Manuel J Rodríguez, Jordi Alberch, Mercè Masana

## Abstract

The alterations in the basal ganglia circuitry associated with motor symptoms in Huntington’s Disease (HD) have been extensively investigated. Yet, the specific contribution of the direct and indirect striatal output pathways from the dorsolateral (DLS) and dorsomedial striatum (DMS) to the motor dysfunction is still not fully understood. Here, using the symptomatic R6/1 male mouse model of HD, strong functional connectivity alterations between DMS and DLS regions with the rest of brain were observed by fMRI, particularly pronounced in the DLS. Then, we systematically evaluated how the selective optogenetic stimulation of the direct and indirect pathways from DLS and DMS influences locomotion, exploratory behavior, and motor learning. In wild type (WT) mice, optogenetic stimulation of the direct pathway from DLS and the indirect pathway from DMS elicited subtle locomotor enhancements, while exploratory behavior remained unaltered. Additionally, stimulation of the indirect pathway from DLS improved the performance in the accelerated rotarod task. In contrast, in HD mice, optogenetic stimulation of the distinct striatal pathways did not modulate these behaviors. Overall, this study points to deficits in the integration of neuronal activity in HD mice, while it contributes to deeper understanding of the complexity of motor control by the diverse striatal subcircuits.

## INTRODUCTION

The dysregulation of basal ganglia circuitry stands as the primary cause behind the manifestation of motor abnormalities in Huntington’s disease (HD). HD is a progressive neurodegenerative disorder that courses with a triad of motor, cognitive and psychiatric symptoms. It is caused by a polyglutamine expansion in the huntingtin gene ^1^, that leads to the degeneration of GABAergic medium-sized spiny neurons (MSNs) of the caudate and putamen nuclei (striatum), the central hub of basal ganglia circuitry.

A complex and coordinated activity of distinct striatal subcircuits modulates locomotion and motor learning processes ^2–5^. Dorsomedial striatum (DMS) and dorsolateral striatum (DLS) subcircuits are based on the clear segregation of cortical inputs along the mediolateral and dorsoventral axes of the dorsal striatum ^6–8^. For instance, cingulate, orbitofrontal and medial prefrontal cortices project mainly to the DMS, while motor and somatosensorial cortices project preferentially to the DLS ^7,8^. Furthermore, the medium-sized spiny neurons (MSNs) from the striatum transmit information through two main pathways—direct and indirect—each projecting to specific output nuclei. Particularly, MSNs from the direct pathway (which express dopamine D1 receptors) project to the internal globus pallidum (GPi) and the substantia nigra reticulata (SNr); while MSNs from the indirect pathway (which express dopamine D2 and A2a receptors) project to external globus pallidum (GPe) ^9^. Notably, both pathways project to their specific output nuclei with segregated mediolateral connections ^10,11^.

However, despite our extensive knowledge of the basal ganglia circuitry under healthy conditions, the involvement of these distinct subcircuits in HD remains limited. In individuals with HD, neurodegeneration primarily occurs in the MSNs of the indirect pathway, with MSNs from the direct pathway being altered later in the course of the disease. In mouse models of HD, impaired motor skill learning linked to cortico-striatal alterations has been extensively described ^12,13^, and alterations in MSNs physiology are observed prior to or in the absence of cell death, affecting both the direct and indirect pathways ^14–16^. Remarkably, cortical afferents to the striatum are important for motor learning ^3^, and selective optogenetic stimulation of M2 cortex-DLS projection restores motor learning and coordination in the R6/1 mouse model of HD ^17^. Therefore, better comprehension of the role of the distinct striatal subcircuits in motor behavior and, specifically, their involvement in HD is needed to further understand motor control and to design effective therapeutic strategies.

Therefore, this study seeks to unveil specific alterations in the distinct basal ganglia subcircuits in HD and its functional implications on locomotion and motor learning. First, we aim to decipher specific alterations in DLS and DMS functional connectivity in HD, by employing *in vivo* resting-state functional MRI (rs-fMRI) techniques using the R6/1 mouse model. Moreover, we evaluate the behavioral consequences of selective optogenetic stimulation of the direct and indirect pathways from DLS and DMS in wild type (WT) and HD mice.

## RESULTS

### Prominent brain functional connectivity deficits from DLS compared to DMS in symptomatic HD mice

Aiming to understand if DLS and DMS are differentially affected in HD mice, we evaluated functional connectivity from DLS and DMS striatum in rs-fMRI using seed-based analysis (Figure 1). First, we segmented the striatum in the MRI atlas template that we have been previously using ^17,18^ into lateral and medial part. To do so, we evaluated anatomical projections maps ^7,8^, and specifically focused in motor and somatosensorial cortex as reference cortices projecting mainly to DLS; and medial prefrontal cortex and cingulate cortex, as reference cortices projecting to the DMS. Based on these projections, we updated the template and analyzed the functional connectivity of both the DLS and the DMS with the rest of the brain (Figure 1A).

**Figure 1.**
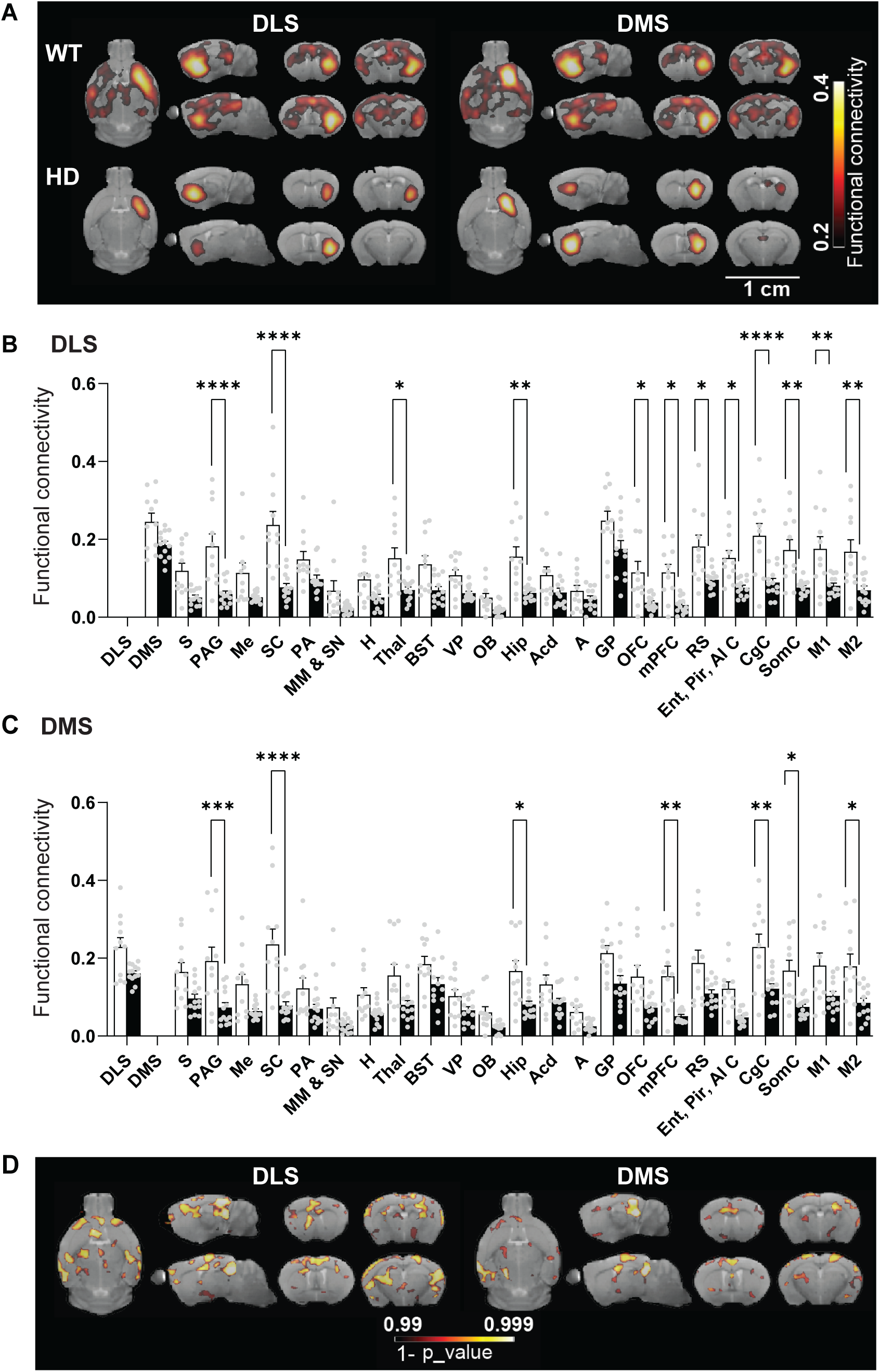
Seed-based functional connectivity using rs-fMRI from DLS and DMS in ∼20-weeks-old WT and HD mice. (A) Average functional connectivity maps from left DLS and left DMS with the whole brain in WT and symptomatic R6/1 mice. Color maps represent the average correlation value when greater than 0.2. (B-C) Mean functional connectivity between (B) left DLS and (C) left DMS with all brain areas from the left hemisphere analyzed are represented. Each point represents data from an individual mouse. Two-way ANOVA with genotype and brain region as factors was performed, followed by Bonferroni post-hoc comparisons test. (D) p-value maps show significant differences voxel-wise between genotypes for DLS and DMS seeds, when smaller than p<0.01. Color code indicates 1 - p_value. Data are represented as mean ± SEM. Number of mice per group: WT n = 11 and HD n = 13 mice. *p<0.05, **p<0.01, ***p<0.001, ****p<0.0001 HD versus WT. Abbreviations: Amygdala (A); Nucleus accumbens (Acd); Bed nucleus of the stria terminalis (BST); Cingulate cortex (CgC); Dorsomedial Striatum (DMS); Dorsolateral Striatum (DLS); Entorhinal, Piriform and Agranular insular cortex (Ent, Pir, AI); Globus pallidus (GP); Hippocampus (Hip); Hypothalamus (H); Medial mammillary nucleus and Substantia nigra (MM & SN); Medial prefrontal cortex (mPFC); Mesencephalic nucleus (Me); Olfactory bulb (OB); Orbitofrontal cortex (OFC); Periaqueductal gray (PAG); Preoptic area (PA); Primary motor (M1) cortex; Secondary motor (M2) cortex; Retrosplenial cortex (RS); Secondary somatosensory cortex (Som); Striatum (Str); Superior colliculus (SC); Thalamus (Thal); Ventral pallidum (VP).

As expected, HD mice showed lower functional connectivity than WT mice, measured by mean correlation value between each pair of regions, from both DLS and DMS. Moreover, the data supports the idea that functional connectivity from DLS is more strongly impaired than from DMS. In one hand, seed-based analysis (Figure 1A,B) from the left DLS reported significant genotype effect (F(1,22) = 15.2; p=0.0008), brain region effect (F(23,506) = 34.52, p<0.0001), and genotype/brain region interaction (F (23, 506) = 4.309, p<0.001), as shown by two-way ANOVA. Compared to WT, left DLS of HD mice showed significantly reduced functional connectivity with left periaqueductal gray, superior colliculus, thalamus, hippocampus, orbitofrontal cortex, medial prefrontal cortex, retrosplenial cortex, entorhinal-piriform-insular cortex, cingulate cortex, somatosensory cortex, primary (M1) and secondary motor cortices (M2), as shown by Bonferroni post-hoc test. In the other hand, seed-based analysis from the DMS (Figure 1A,C) showed genotype effect (F(1,22) = 12.3; p=0.002), brain region effect (F(23,506) = 25.63, p<0.0001) and genotype/brain region interaction effect (F(23,506) = 3.204, p<0.0001), analyzed by two-way ANOVA. Bonferroni post-hoc test showed that functional connectivity from DMS of HD mice was significantly reduced with left periaqueductal gray, superior colliculus, hippocampus, medial prefrontal cortex, cingulate cortex, somatosensory cortex and M2 cortex, compared to WT.

Additionally, to further assess the genotype differences in the DLS and the DMS connectivity maps, a voxel-wise analysis was performed to precisely identify location and number of voxels with statistical differences in mean correlation values between HD and WT mice (Figure 1D). Out of 63593 voxels analyzed, the DLS exhibited significant differences in mean correlation values among genotypes in 203 mm³ (25378 voxels), while the DMS revealed significant differences in a total of 155 mm³ (19461 voxels). Overall, our results highlight functional connectivity alterations in both DLS and DMS of symptomatic HD mice, with major deficits from the DLS compared to DMS.

### Light stimulation pattern differently modulates motor behavior in wild type mice

To further understand how the distinct striatal subcircuits modulate motor activity, we took advantage of available optogenetic tools and mice that express Cre under the D1 and A2a-promoter. Expression of Cre in direct and indirect pathways of the striatum was confirmed by crossing the D1-Cre and A2a-Cre mice with flox-TdTomato and evaluating their pathway-specific expression (Figure S1A). Then, we transduced mice with an AAV expressing ChR2 (or its control YFP) under double-floxed inverse open reading frame (DIO) system in DLS to achieve opsin expression either in D1 or A2a expressing neurons. Fiber-optic cannulas were implanted in the same coordinates. We tested two light stimulation protocols based on previous studies: constant illumination ^19^ and 10 Hz light stimulation ^17^. Constant illumination was able to induce contralateral circling in D1 DLS WT mice and contralateral circling in A2A DLS WT mice during stimulation (Figure S1B), while 10 Hz stimulation was not able to induce clear behavioral effects (Figure S1C). Thus, constant illumination pattern was selected as stimulation protocol for subsequent experiments.

### Distinct locomotor responses are induced by optogenetic stimulation of the direct or indirect pathways from DLS and DMS in WT and HD mice

We aimed to understand how selective activation of direct and indirect pathways in the DLS or DMS could show differential motor responses in WT and the R6/1 mouse model of HD. First, we crossed D1-Cre and A2a-Cre mice with R6/1 mice to obtain D1-WT, D1-HD, A2a-WT and A2a-HD mice. AAV-YFP or AAV-ChR2 was injected in the DLS or DMS of these mouse groups. Four weeks later, we evaluated how specific optogenetic activation of DMS and DLS striatal subcircuits affects motor behaviors in both WT and HD mice. The experimental procedure is detailed in Figure 2. Mice were optogenetically stimulated once per week, during three consecutive weeks, while mice performed an OF task (Figure 2B). The day after the second session of OF, mice performed the ARR to assess motor learning. DMS and DLS subcircuits were illuminated in a series of 10 trials during the OF, each trial alternated 30 s during which the laser was ON (constant illumination), followed by 60 s period during which the laser was OFF (Figure 2B). Validation of correct viral expression and cannula implantation was assessed by immunofluorescence (Figure 2C).

**Figure 2.**
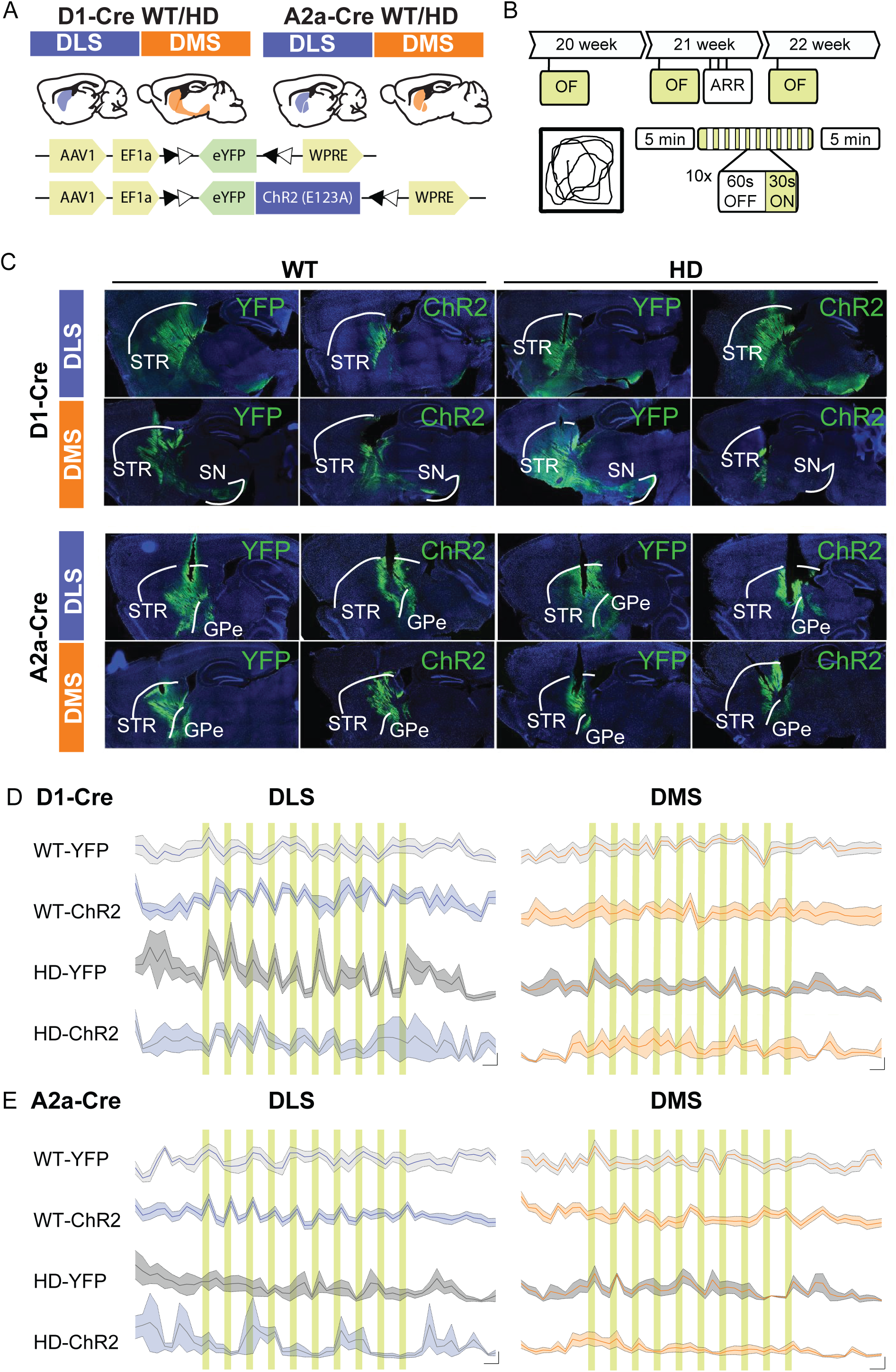
Experimental protocol for the optogenetic stimulation of the direct or indirect pathways from DLS and DMS in WT and HD mice. (A) Schematic representation of the experimental mouse groups and AAV constructs. D1-Cre and A2a-Cre mice were crossed with R6/1 mice to obtain D1-WT, D1-HD, A2a-WT and A2a-HD mice. Then, AAV-YFP or AAV-ChR2 constructs were injected in the DLS or DMS of these mouse groups. (B) Optogenetic stimulation was performed during the three OF procedures at 20, 21 and 22 weeks of mouse age. The accelerating rotarod (ARR) task was performed after the 2nd stimulation day. Optogenetic stimulation started five minutes after the mice was placed in the OF and consisted of 10 trials of laser illumination and each trial alternated a 30 s period in which laser was ON [Constant illumination, ∼5 mW] and 60 s period in which laser was OFF. Mice were left in the OF for an additional five minutes after the stimulation train. (C) Representative immunofluorescence images of AAV-ChR2 and AAV-YFP constructs expression (green), DAPI (blue) and cannula location at DLS or DMS of D1 WT, D1 HD, A2a WT and A2a HD mice. (D-E) Average distance travelled (cm) per min during the first OF session are represented for WT YFP, WT ChR2, HD YFP and HD ChR2 expressed in (D) direct pathway of the DLS (left panel) and DMS (right panel) and (E) indirect pathway from the DLS (left panel) and the DMS (right panel). The green shadow indicates the ten consecutive periods of 30 s in which laser was ON. Data are represented as mean ± SEM. Number of mice per group: Drd1-DLS-WT-YFP n = 7, Drd1-DLS-WT-ChR2 n = 5, Drd1-DLS-HD-YFP n = 5, Drd1-DLS-HD-ChR2 n = 4, Drd1-DMS-WT-YFP n = 7, Drd1-DMS-WT-ChR2 n = 5, Drd1-DMS-HD-YFP n = 5 and Drd1-DMS-HD-ChR2 n = 3; A2a-DLS-WT-YFP n = 7, A2a-DLS-WT-ChR2 n = 12, A2a-DLS-HD-YFP n = 4, A2a-DLS-HD-ChR2 n = 3, A2a-DMS-WT-YFP n = 10, A2a-DMS-WT-ChR2 n = 12, A2a-DMS-HD-YFP n = 4 and A2a-DMS-HD-ChR2 n = 6. Abbreviations: Globus Pallidus pars externa (GPe), Substantia nigra (SN) and Striatum (STR).

In bilaterally stimulated mice, the stimulation induced changes in locomotor activity throughout the OF. Figure 2D-E shows the precise changes in locomotor activity every 30s in D1 DLS, D1 DMS, A2a DLS and A2a DMS. Each panel shows the effects of WT and HD mice expressing either ChR2 or YFP. Our data shows a clear light-dependent locomotor pattern in mice expressing both ChR2 and YFP, indicating that light *per se* might have an influence on these locomotor changes. Moreover, the profile of these changes decreased over time, suggesting certain habituation to the light and the open field.

Thus, to avoid light-dependent but not ChR2-dependent effects, we decided to only compare the effects on locomotion the 5 min before (PRE) and the 5 min after (POST) stimulation (Figure 3). During the 5 min PRE-stimulation, there were no significant differences in locomotor activity between WT-YFP, WT-CHR2, HD-YFP and HD-CHR2 groups over the three OF sessions for all the pathways analyzed except in mice in which optogenetic stimulation was performed in the indirect pathway from the DMS (Table1). In this group, Bonferroni main group post-hoc comparisons showed that optogenetic stimulation increased locomotor activity in WT-CHR2 compared to WT-YPF.

**Figure 3.**
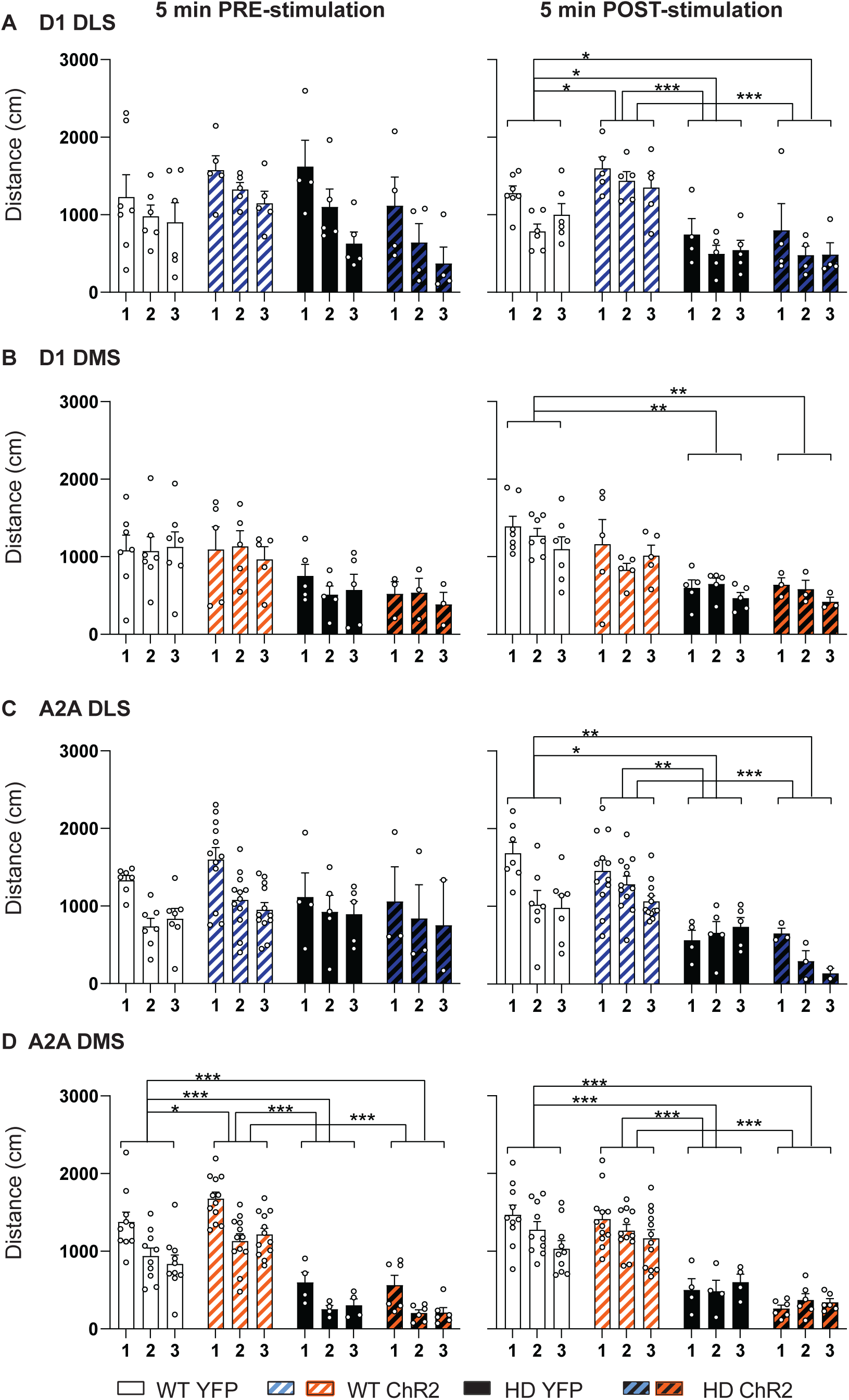
Locomotor activity induced by optogenetic stimulation of the direct (D1) or indirect (A2a) pathways from DLS and DMS in WT and HD mice, over three open field sessions. The effects of ChR2 or YFP expressed in WT and HD mice on total distance travelled was compared before (5 minutes PRE, left panels) and after (5 minutes POST, right panels) the optogenetic stimulation for (A) direct pathway of the DLS (D1 DLS), (B) direct pathway from the DMS (D1 DMS), (C) indirect pathway from the DLS (A2a DLS) and (D) indirect pathway from the DMS (A2a DMS). A two-way ANOVA with AAV construct and genotype group as factors was performed, followed by main group Bonferroni post-hoc comparisons test. Each point represents data from an individual mouse and data are represented as mean ± SEM. Number of mice per group: Drd1-DLS-WT-YFP n = 7, Drd1-DLS-WT-ChR2 n = 5, Drd1-DLS-HD-YFP n = 6, Drd1-DLS-HD-ChR2 n = 4, Drd1-DMS-WT-YFP n = 7, Drd1-DMS-WT-ChR2 n = 5, Drd1-DMS-HD-YFP n = 5 and Drd1-DMS-HD-ChR2 n = 3; A2a-DLS-WT-YFP n = 7, A2a-DLS-WT-ChR2 n = 12, A2a-DLS-HD-YFP n = 5, A2a-DLS-HD-ChR2 n = 3, A2a-DMS-WT-YFP n = 10, A2a-DMS-WT-ChR2 n = 12, A2a-DMS-HD-YFP n = 4 and A2a-DMS-HD-ChR2 n = 6. *p<0.05, **p<0.01, ***p<0.001.

Conversely, in the 5 min POST-stimulation, we clearly observed group effects in all conditions, as summarized in Table 1. Bonferroni main group post-hoc comparisons showed significantly reduced locomotion in HD mice in all conditions, compared to WT YFP. Optogenetic stimulation was unable to modulate locomotion in HD mice in any of the striatal subcircuits tested. Of note, locomotion was significantly induced in WT-CHR2 mice compared to WT-YPF mice when the direct pathway from the DLS was stimulated (Figure 3A).

**TABLE 1.**

| Group | Factor | Locomotion |  | Rearing time |  |
| --- | --- | --- | --- | --- | --- |
|  |  | F(DFn, DFd) | P value | F(DFn, DFd) | P value |
| D1 | Day | F (2, 30) = 18.94 | **<0.0001\*** | F (2, 30) = 0.058 | 0.9438 |
| DLS | Group | F (3, 18) = 1.532 | 0.2404 | F (3, 18) = 3.958 | **0.0249\*** |
| PRE | Day/Group | F (6, 30) = 0.855 | 0.5384 | F (6, 30) = 0.296 | 0.9342 |
| D1 | Day | F (2, 30) = 5.225 | **0.0113\*** | F (2, 30) = 4.241 | **0.0239\*** |
| DLS | Group | F (3, 18) = 16.62 | **<0.0001\*** | F (3, 18) = 7.610 | **0.0017\*** |
| POST | Day/Group | F (6, 30) = 0.343 | 0.9086 | F (6, 30) = 0.932 | 0.4867 |
| D1 | Day | F (2, 32) = 0.897 | 0.4180 | F (2, 32) = 2.046 | 0.1459 |
| DMS | Group | F (3, 16) = 2.644 | 0.0846 | F (3, 16) = 4.882 | **0.0135\*** |
| PRE | Day/Group | F (6, 32) = 0.699 | 0.6521 | F (6, 32) = 1.752 | 0.1411 |
| D1 | Day | F (2, 32) = 2.740 | 0.0797 | F (2, 32) = 3.230 | 0.0527 |
| DMS | Group | F (3, 16) = 9.375 | **0.0008\*** | F (3, 16) = 5.995 | **0.0061\*** |
| POST | Day/Group | F (6, 32) = 0.797 | 0.5795 | F (6, 32) = 1.673 | 0.1599 |
| A2A | Day | F (2, 44) = 8.768 | **0.0006\*** | F (2, 44) = 0.471 | 0.6274 |
| DLS | Group | F (3, 23) = 0.927 | 0.4433 | F (3, 23) = 3.515 | **0.0312\*** |
| PRE | Day/Group | F (6, 44) = 1.143 | 0.3538 | F (6, 44) = 1.472 | 0.2101 |
| A2A | Day | F (2, 44) = 8.962 | **0.0005\*** | F (2, 44) = 0.071 | 0.9316 |
| DLS | Group | F (3, 23) = 11.26 | **<0.0001\*** | F (3, 23) = 8.979 | **0.0004\*** |
| POST | Day/Group | F (6, 44) = 2.732 | **0.0241\*** | F (6, 44) = 0.099 | 0.9962 |
| A2A | Day | F (2, 56) = 21.17 | **<0.0001\*** | F (2, 56) = 1.800 | 0.1748 |
| DMS | Group | F (3, 28) = 45.28 | **<0.0001\*** | F (3, 28) = 10.45 | **<0.0001\*** |
| PRE | Day/Group | F (6, 56) = 0.480 | 0.8208 | F (6, 56) = 0.387 | 0.8840 |
| A2A | Day | F (2, 56) = 1.837 | 0.1688 | F (2, 56) = 0.389 | 0.6794 |
| DMS | Group | F (3, 28) = 29.55 | **<0.0001\*** | F (3, 28) = 15.45 | **<0.0001\*** |
| POST | Day/Group | F (6, 56) = 2.178 | 0.0586 | F (6, 56) = 0.595 | 0.7332 |

Summary of two-way ANOVA statistical analysis of locomotion and rearing time before (PRE) and after (POST) optogenetic stimulation for each striatal subcircuit group: D1 DLS, D1 DMS, A2a DLS and A2a DMS. Four conditions were compared over three stimulation sessions in the OF for each subcircuit analyzed: WT YFP, WT CHR2, HD YFP and HD ChR2.

### Exploratory behavior is not modulated by optogenetic stimulation of the distinct striatal output pathways in either WT or HD mice

We evaluated the effects of the optogenetic stimulation on the rearing time, as optogenetic stimulation of M2 cortex-DLS circuit was able to modulate exploratory activity in HD mice ^17^ (Figure 4). We compared the effect of optogenetic stimulation on total rearing time before (PRE) and after (POST) the stimulation and over the 3 sessions in D1 DLS, D1 DMS, A2a DLS and A2a DMS striatal subcircuits tested.

**Figure 4.**
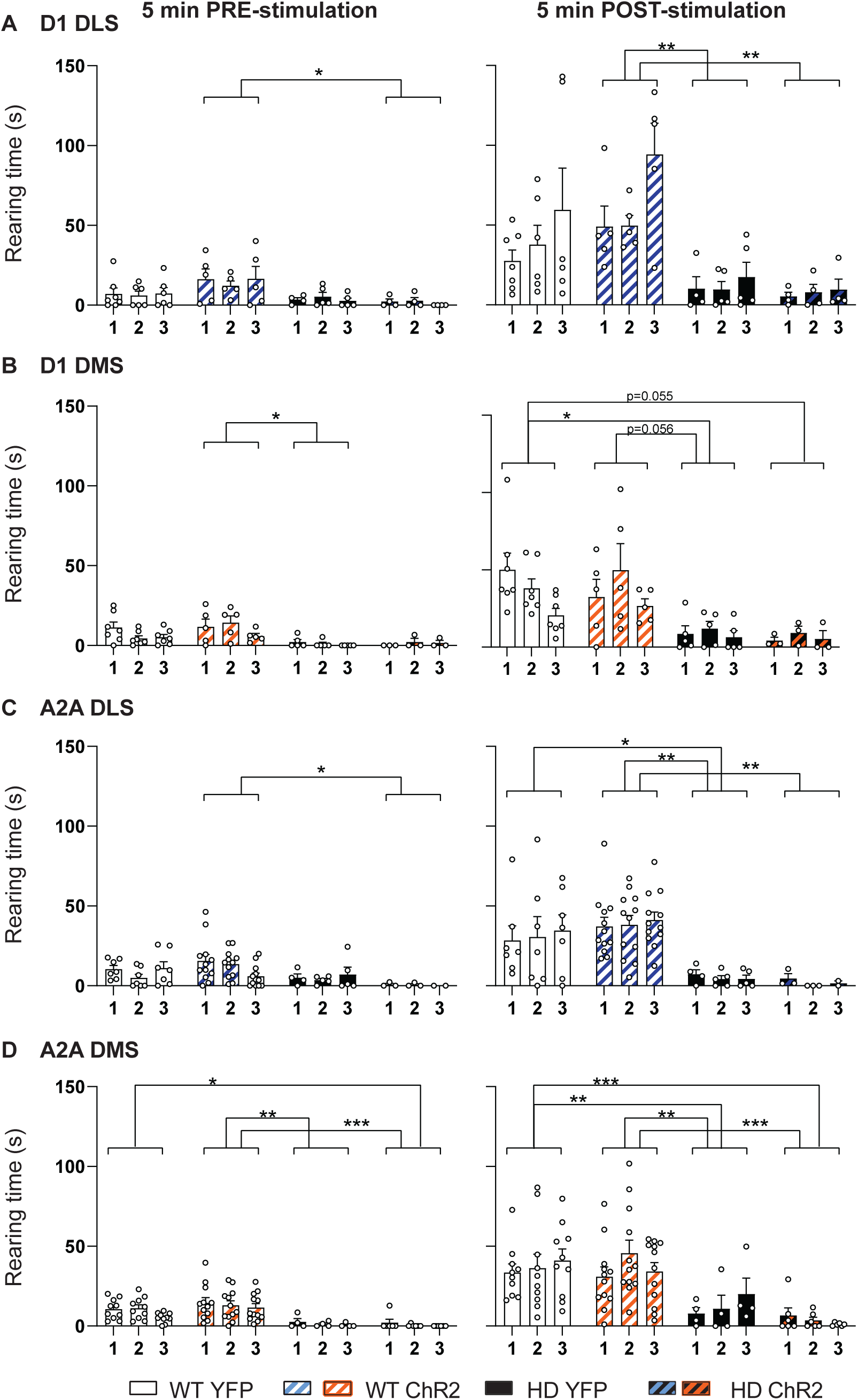
Exploratory behavior induced by optogenetic stimulation of the direct (D1) or indirect (A2a) pathways from DLS and DMS in WT and HD mice, over three open field sessions. The effects of ChR2 or YFP expressed in WT and HD mice on total rearing time was compared before (5 minutes PRE, left panels) and after (5 minutes POST, right panels) the optogenetic stimulation for (A) direct pathway of the DLS (D1 DLS), (B) direct pathway from the DMS (D1 DMS), (C) indirect pathway from the DLS (A2a DLS) and (D) indirect pathway from the DMS (A2a DMS). A two-way ANOVA with AAV construct and genotype group as factors was performed, followed by main group Bonferroni post-hoc comparisons test. Each point represents data from an individual mouse and data are represented as mean ± SEM. Number of mice per group: Drd1-DLS-WT-YFP n = 7, Drd1-DLS-WT-ChR2 n = 5, Drd1-DLS-HD-YFP n = 6, Drd1-DLS-HD-ChR2 n = 4, Drd1-DMS-WT-YFP n = 7, Drd1-DMS-WT-ChR2 n = 5, Drd1-DMS-HD-YFP n = 5 and Drd1-DMS-HD-ChR2 n = 3; A2a-DLS-WT-YFP n = 7, A2a-DLS-WT-ChR2 n = 12, A2a-DLS-HD-YFP n = 5, A2a-DLS-HD-ChR2 n = 3, A2a-DMS-WT-YFP n = 10, A2a-DMS-WT-ChR2 n = 12, A2a-DMS-HD-YFP n = 4 and A2a-DMS-HD-ChR2 n = 6. *p<0.05, **p<0.01, ***p<0.001.

Two-way ANOVA with group and day as factors showed that there are significant group effects for all experimental groups tested, as detailed in Table 1. However, Bonferroni main group comparisons post hoc test showed significant genotype interactions, but not significant effect of CHR2 expressing groups compared to YFP expressing groups, indicating that the optogenetic stimulation of the diverse striatal subcircuits was not able to modulate rearing time in neither WT nor HD mice.

### Optogenetic stimulation of the indirect pathway from the DLS increases motor learning in WT mice but not in HD mice

We further explored the effects of the distinct striatal output subcircuit stimulation on motor learning. Motor learning was assessed by measuring the latency to fall from the ARR, as previously described ^12,13,17^. As expected, HD mice showed reduced latency to fall compared to WT in all groups tested (Figure 5). Notably, optogenetically stimulated WT mice increased the latency to fall from the rod when the stimulation was performed in the indirect pathway from DLS (Figure 5C). Moreover, we quantified the area under the curve (AUC) for each experimental group (Figure 5, right panel). One-way ANOVA showed significant group effects in all tested sub-circuits (D1 DLS: (F(3,15) = 14.5, p=0.0001); D1 DMS: (F(3,16) = 11.6, p=0,0003); A2a DLS: (F(3,23) = 22.75, p<0,0001) and A2A DMS: (F(3,26) = 8.422, p=0,0004) ). Bonferroni post-hoc test highlight that A2A DLS optogenetically stimulated WT mice (WT CHR2) increase motor learning performance compared to control (WT YFP) (p=0,05), while all other significant group comparisons reflect genotype differences in all subcircuits evaluated. As for locomotion and exploratory behavior, optogenetic stimulation did not modulate motor learning in any HD mouse group.

**Figure 5.**
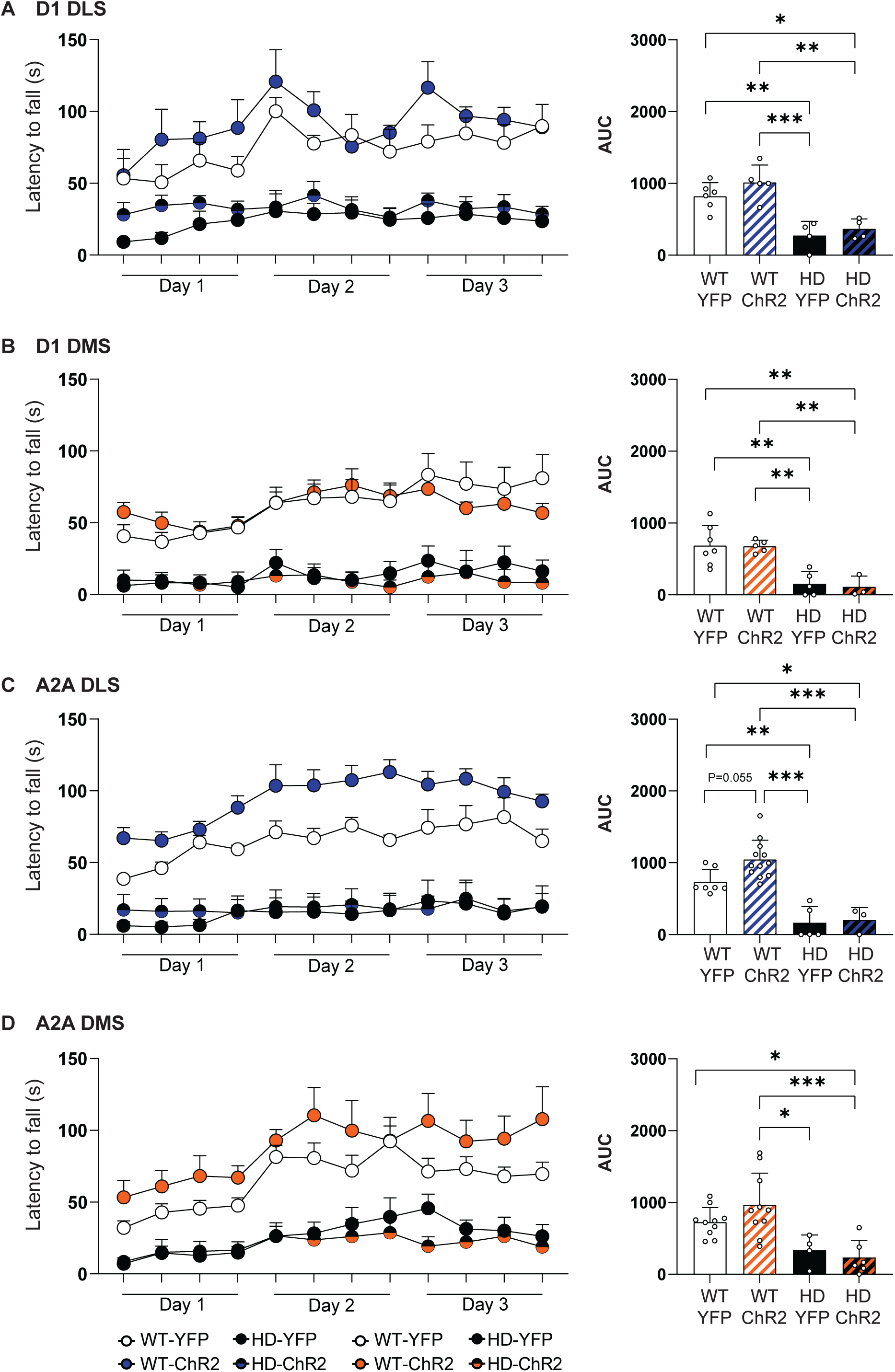
Effects of optogenetic stimulation of the direct (D1) or indirect (A2a) pathways from DLS and DMS in WT and HD mice on motor learning. Latency to fall in the ARR was measured after two sessions of optogenetic stimulation of (A) direct pathway of the DLS (D1 DLS), (B) direct pathway from the DMS (D1 DMS), (C) indirect pathway from the DLS (A2a DLS) and (D) indirect pathway from the DMS (A2a DMS). Additionally, the area under the curve (AUC) was calculated. AUC data were analyzed by one-way ANOVA, followed by Bonferroni’s mean group multiple comparison test as post-hoc. Each point represents data from an individual mouse and data are represented as mean ± SEM. Number of mice per group: Drd1-DLS-WT-YFP n = 6, Drd1-DLS-WT-ChR2 n = 5, Drd1-DLS-HD-YFP n = 4, Drd1-DLS-HD-ChR2 n = 4, Drd1-DMS-WT-YFP n = 7, Drd1-DMS-WT-ChR2 n = 5, Drd1-DMS-HD-YFP n = 5 and Drd1-DMS-HD-ChR2 n = 3; A2a-DLS-WT-YFP n = 7, A2a-DLS-WT-ChR2 n = 12, A2a-DLS-HD-YFP n = 5, A2a-DLS-HD-ChR2 n = 3, A2a-DMS-WT-YFP n = 10, A2a-DMS-WT-ChR2 n = 10, A2a-DMS-HD-YFP n = 4 and A2a-DMS-HD-ChR2 n = 6. *p<0.05, **p<0.01, ***p<0.001.

## DISCUSSION

Alterations in the basal ganglia circuitry are responsible for the motor symptoms in HD. However, the contribution of the distinct striatal subcircuits to HD phenotype has not been thoroughly explored. Here, we evaluate the functional connectivity of DLS and DMS with the rest of the brain. Our data reveal strong functional connectivity alterations in HD mice of both striatal regions, although functional connectivity from the DLS shows alterations with more brain regions, suggesting a prominent alteration in this striatal region, at least at symptomatic stages. Moreover, we systematically evaluated the effects on locomotion, exploratory behavior and motor learning induced by the selective optogenetic stimulation of the direct and indirect pathway from DLS and DMS, in WT and HD mice. Surprisingly, optogenetic stimulation of striatal projection neurons from the distinct subcircuits subtly modulated motor responses in WT mice, while was unable to modify any motor phenotype in HD mice.

Our MRI data shows that both DMS and DLS regions from our R6/1 mouse model display prominent functional connectivity deficits with cortical regions in symptomatic mice, in line with previous publications ^17,20,21^. Particularly, cortical afferents from M2, somatosensory and mPFC might have a prominent role in the symptomatic R6/1 mouse model of HD, as their connectivity is altered with both striatal subregions. Indeed, in HD patients, specific alterations in the cortex are associated to the phenotypic heterogeneity of symptoms ^22,23^, and differential dynamics of cortical degeneration has also been described ^24^. This is also in line with the differential volume loss observed between caudate and putamen, which also show differential loss rate ^24,25^. Thus, DMS and DLS subcircuit alterations might occur also at different rates. Yet, further longitudinal studies would be needed to address this point.

Additionally, our data shows strong functional connectivity alterations from both DLS and DMS with the superior colliculus and its direct output region the periaqueductal gray. Of note, the superior colliculus receive collateral inputs from M2 and cingulate cortex and is involved in the control of saccadic movements ^26,27^, which are altered from presymptomatic stages in people with HD ^28,29^. Moreover, alterations in M2-superior colliculus circuitry has been recently described and associated to sensory-induced behavioral alterations in HD mice ^30^. Thus, the strong functional connectivity alterations from both DMS and DLS to the superior colliculus and periaqueductal gray support the idea that these brain regions might have a role in the basal ganglia alterations in HD. To delve into the contributions of the distinct striatal subcircuits to motor control, we employed optogenetic techniques to selectively stimulate D1 or A2a expressing neurons in either the DLS or DMS. While unilateral optogenetic stimulation reliably induced rotations, bilateral stimulation yielded more subtle locomotion effects. These findings challenged the prevailing notion that optogenetic activation of D1 or A2a expressing cells strictly elicits or inhibits movement, as previously suggested ^19^. Such assumptions have been supported by lesion studies demonstrating that specific ablation of the direct pathway reduces locomotion, while ablation of the direct pathway increases it ^2^. Notably, these effects were confined in the DMS ^2^, suggesting a prominent role of DMS MSNs in ambulatory behavior. However, recent literature has refined this view, pointing that the observed effects on locomotion might reflect changes in reinforcing or punishment processes, rather than direct motor control ^31,32^. Indeed, recent research indicates that motor suppression can be induced by increasing stimulation light intensity in A2A-MSNs, an effect attributed to the engagement of sufficient striatal collateral inhibition rather than indirect pathway activation ^32^. Therefore, it seems that changes in the behavioral context, MSN pattern of activity and the extent of striatal volume stimulated/inhibited or lesioned might contribute to the heterogeneous behavioral responses observed, an observation previously pointed by others ^33^.

Regarding motor learning, our findings reveal that prior optogenetic stimulation of A2a-expressing neurons in the DLS enhances performance on the accelerated rotarod task in WT mice. This aligns with previous research suggesting that extensive training induces long-term potentiation (LTP) onto A2a-expressing neurons from the DLS ^5^. However, the low number of mice in the present study does not allow to exclude the involvement of the other subcircuits in motor learning. Indeed, several authors have shown that the acquisition of a skill requires a complex re-organization of MSNs neuronal activity in both DLS and DMS ^3–5^. Moreover, ablation of the direct pathway in the DLS, but not the DMS, impairs the acquisition and performance of the accelerating rotarod task, while ablation of the indirect pathway in the DMS delays learning but does not affect performance once mice already learned ^2^. Furthermore, motor learning increased activity in both DLS and DMS inputs in the early stages of learning, although involving pre-synaptic potentiation of cortico-striatal function ^3^. Therefore, complementary approaches able to dynamically evaluate neuronal activity simultaneously in the different cortico-striatal subcircuits during the accelerated rotarod task are still required to fully understand the contribution of each subcircuit to motor learning.

Motor learning deficits and reduced exploratory behavior are core symptoms present in HD models ^12,13,17,34,35^. However, inducing neuronal activity in the distinct striatal subcircuits with optogenetics was not sufficient to modulate motor responses in HD mice, despite unilateral stimulation was able to induce rotational behavior also in HD mice (data not shown). On the one hand, these results support the idea that striatal output pathways will only modulate locomotion when engaging striatal collaterals within the striatum ^32^. This is in line with the aberrant communication between direct and indirect pathways within the striatum in different HD mouse models ^14–16,36^, which is accompanied by a significant decrease in the amplitude of optogenetically-induced GABA responses in the SNr ^14^. On the other hand, optogenetic stimulation of cortical-afferents to the DLS ameliorates motor learning and exploratory behavior deficits in the R6/1 mice ^17^, while direct stimulation of MSNs in the DLS (present results) does not impact the learning nor performance, neither exploratory activity in R6/1 mice at same symptomatic stage. Accordingly, it has been described that motor learning requires cortical afferent reorganization in the striatum ^3^. Therefore, acting on cortico-striatal circuitry rather than directly in the striatum, seems a better strategy to ameliorate HD motor symptoms.

In summary, our results highlight the prominent functional connectivity alterations in DLS compared to DMS in HD. Moreover, optogenetic stimulation of direct and indirect pathways from DLS and DMS induces subtle locomotor and motor learning responses in healthy mice, while the lack of responses by optogenetic stimulation in HD mice points to deficits in the integration of neuronal activity in the HD striatum. Thus, our results favor the idea that the orchestrated activity of the distinct striatal subcircuits is more intricate than expected and that strategies involving cortico-striatal circuits, rather than direct striatal stimulation might be a better therapeutic strategy for HD. Yet, further understanding the complexity of motor control is still needed to better design therapeutic strategies for HD.

## MATERIALS AND METHODS

### Animals

The transgenic R6/1 mice were used as mouse model of HD. R6/1 mouse express the exon-1 of mutant huntingtin with ∼145 CAG repeats and were originally acquired from The Jackson Laboratory (Bar Harbor, ME; (B6CBA-Tg(HDexon1)61Gpb/1J; RRID:IMSR_JAX:002809) and maintained on a B6CBA background by breeding transgenic male mice with C57BL/6J x CBA/J F1 females. Heterozygous R6/1 male mice were crossed with heterozygous Adora2a-Cre+/- ^37^ and heterozygous Drd1a-Cre+/- female (129S6.FVB(B6)-Tg(Drd1a-cre)AGsc/KndlJ, The Jackson Laboratory) to obtain WT-Adora2a-Cre+ (A2a-WT mice), R6/1-Adora2a-Cre+ (A2a-HD mice), R6/1-Drd1-Cre+ (D1-HD mice) and WT-Drd1a-Cre+ (D1-WT mice), respectively. Genotypes were determined by polymerase chain reaction from ear biopsy and data were recorded for analysis by microchip mouse number. Mice were housed in groups of mixed genotypes under a 12:12 hr light/dark cycle with access to water and food ad libitum, in a room kept at 19-22 °C and 40-60% humidity. A maximum of 5 mice were housed together in a cage containing sawdust and enrichted with wooden bricks and shredded paper. Experiments were conducted using 16∼22 weeks-old male mice. All animal procedures were conducted in accordance with the Spanish RD 53/2013 and European 2010/63/UE regulations for the care and use of laboratory animals and approved by the animal experimentation Ethics Committee of the Universitat de Barcelona (274/18) and Generalitat de Catalunya (10101). This study was in compliance with the ARRIVE guidelines.

### MRI acquisition

A 7.0T BioSpec 70/30 horizontal animal scanner (Bruker BioSpin, Ettlingen, Germany), equipped with an actively covered gradient structure (400 mT/m, 12 cm inner diameter) was used to scan WT mice and R6/1 mice at 17∼20 weeks of age. Mice were placed in a supine position in a Pexiglas holder and were fixed in a nose cone using tooth and ear bars and adhesive tape; a combination of anesthetic gases [medetomidine (bolus of 0.3 mg/kg, 0.6 mg/kg/h infusion) and isoflurane (0.5%)].

A 3D-localizer scan was used to ensure the accurate position of the head at the isocenter of the magnet. T2-weighted image was obtained using a RARE sequence [effective TE = 33 ms, TR = 2.3 s, RARE factor = 8, voxel size = 0.08 x 0.08 mm2, slice thickness = 0.5 mm]. rs-fMRI acquisition was performed using an echo planar imaging (EPI) sequence [TR = 2 s, TE = 19.4, voxel size 0.21 x 0.21 mm2, slice thickness = 0.5 mm]; 420 volumes were acquired during 14 min. Atipamezol (Antisedan®, Pfizer) and saline were injected after the imaging session to reverse sedative effects and compensate fluid loss.

### Functional connectivity analysis

Seed-based analysis was performed as previously described ^17,18^ to evaluate functional connectivity of the DLS and DMS with the rest of the brain. Pre-processing of the rs-fMRI data was performed using NiTime (http://nipy.org/nitime) and nilearn (https://nilearn.github.io) and included: slice timing, spatial realignment using SPM8 for motion correction, elastic registration to the T2-weighted volume using ANTs for EPI distortion correction ^38^, detrend, smoothing [full-width half maximum (FWHM) = 0.6 mm], frequency filtering of time series between 0.01 and 0.1 Hz and regression by motion parameters.

Parcellation of the brain was defined according to an MRI-based mouse atlas ^39^ with some modifications. First, the sensorimotor cortex was manually divided into somatosensory cortex (SomC), orbitofrontal cortex (OFC), primary and secondary motor cortex (M1 and M2 respectively), cingulate cortex (CgC), medial prefrontal cortex (mPFC) and retrosplenial cortex (RSC). Then, the striatum was subdivided into the DLS and the DMS. The resulting atlas template was elastically registered to each subject T2-weighted volume using ANTs ^38^ to obtain brain regions parcellation for each animal and then to the corresponding resting-state fMRI images.

Left DLS and left DMS were selected from the automatic parcellation for seed-based analysis. A correlation map describing the connectivity of each seed with the rest of the brain. Extracted average time series in the DLS or the DMS were correlated with the time series of all the voxels within the brain respectively, resulting in connectivity maps containing the value of the correlation between the BOLD signal time series in each voxel with the seed time series. Seed to region connectivity was calculated as the mean value of the correlation map in each region, considering only positive correlations.

### Stereotaxic surgery

Stereotaxic surgery was performed in ∼16-week-old male D1-HD, D1-WT, A2a-HD and A2a-WT mice under isoflurane anesthesia (5% induction and 1.5% maintenance). After fur shaving and scalp cleaned with ethanol and iodine, local anesthesia was applied (Lidocaine 2.5% and Prilocaine 2.5% EMLA, AstraZeneca) and a dose of 2 mg/kg of analgesic Metacam was injected subcutaneously. Small bilateral holes were drilled according to DLS or DMS coordinates from bregma and dura matter: DLS [+0.1 AP, ±2.2 L, -3.0 DV]; DMS [+0.5 AP, ±1.5 L, -3.0 DV] and double-floxed inverted (DIO) recombinant AAV-DIO encoding the channelrhodopsin-2 (ChR2) fused to enhanced yellow fluorescent protein (eYFP) (AAV1-EF1a-DIO-ChR2(H134R)-eYFP; Addgene catalog: #20298-AAV1) or eYFP as control (AAV5-EF1a-DIO-eYFP; Addgene catalog: #27056-AAV5) were injected. 0,5 μL of the AAV construct was injected in each hemisphere using 5 µl Hamilton syringe with a 33-gauge needle at 0.1 µl/min, followed by an additional 5 min period to allow diffusion and avoid reflux. Then, fiber-optic cannulas (MFC_200/240-0.22_3.5mm_ZF1.25_FLT; Doric Lenses) were implanted and fixed during surgery using dental cement (TAB 2000tm, Kerr Dental). All surgeries were performed 4 weeks before experiments were initiated to allow mice recovery and a stable expression of viral constructs. AAV expression and cannula implantation was validated post-mortem by immunofluorescence in sagittal sections of one hemisphere and mice were excluded from the analysis when AAV was not expressed and/or cannula was misplaced.

### Optogenetic stimulation

Blue light was delivered from 473 nm diode-pumped solid-state blue laser (Laserglow) using a custom-made wave-form generator (Arduino) while mice were freely moving in the open field (OF) at 20, 21 and 22-weeks of age. Mice were placed in the OF and after 5 min, light stimulation was performed in a series of 10 trials of laser illumination and each trial alternated a 30 s period in which laser was ON [Cte illumination, ∼5 mW (at tip of each fibre)], followed by a 60 s period in which laser was OFF, based on ^19^, then mice were left in the OF for 5 additional minutes.

### Behavioral assessment

Behavioral tests were performed as previously described ^17^. Animals were habituated to the experimental room for at least 1 hour before testing and all apparatus were cleaned with water and dried between animals. Mice that experienced a seizure just before or during the test were excluded from the corresponding data and not included in the analyses.

#### Open Field (OF)

Optogenetic stimulation was performed in the OF, which consists in a white square arena (40 x 40 x 30 cm3) with dim light (∼20 lux). Mice were left in the center of the apparatus and allowed to freely explore the arena during a total of 25 min (5 min PRE-stimulation, 15 min TRAIN-stimulation (30sON+60OFF), 5 minutes POST-stimulation. Tracking and recording of mice were performed using SMART 3.0 software (Panlab).

#### Accelerating rotarod (ARR)

Motor learning was assessed using the rotarod apparatus at 21 weeks of age. Mice were placed on a motorized horizontal rod (30 mm diameter) with a rotation speed increasing from 4 to 40 rpm over 5 min. The test was performed four times per day during three consecutive days (12 trials in total) and latency to fall from the rod was measured for each trial. Different trials during the same day were separated by 1 hr.

### Immunohistochemistry

Intra-cardiac perfusion was performed with cold PBS and 4% paraformaldehyde solution. Brains were post-fixed in 4% paraformaldehyde for 24 h and dehydrated in a PBS/Sucrose gradient [from 15% (48 h post-mortem) to 30% (72 h post-mortem)] with 0.02% Sodium Azide and kept at 4°C. Sagittal sections were obtained at 30 µm with a vibratome (Leica VT1000 S) and cryopreserved [30% Ethylene glycol, 30% Glycerol, 25% Tris HCl (pH=7.5) and 15% H2O miliQ] at -20°C.

Viral injections were validated using anti-GFP (anti-Green Fluorescent Protein, 1:500, Invitrogen, #11122) as previously described ^17^. Free-floating sections were washed in PBS and permeabilized and blocked for 15 min in PBS containing 0.3% Triton X-100 and 3% Normal Goat Serum (NGS, Pierce Biotechnology). Sections were then washed again in PBS and incubated overnight at 4°C with primary antibodies. The day after, slices were washed three times and then incubated 2 h rocking at RT with the secondary antibody goat anti-rabbit Cy3 (Cy3, 1:200, Jackson ImmunoResearch, West Grove, PA, USA). Brain slices were finally mounted on microscope slides using DAPI Fluoromount-G (SouthernBiotec). Fluorescence signal was detected using a Leica AF600 Fluorescence Microscope and mosaics were captured using a 20x objective with 1.6 magnification.

### Statistical analysis

Results are expressed as mean ± SEM and data from individual mouse is represented by single points when possible. GraphPad Prism version 8.0.0. Software was used for statistical analyses. Statistical analyses included one-way ANOVA or two-way ANOVA analysis and followed by Bonferroni post-hoc test, with the factors used indicated in the results section and/or figure legends. Values were considered as statistically significant when p<0.05. Differences between the seed-based connectivity maps of the DMS and the DLS were further evaluated voxel-wise using the Randomise method implemented in FSL ^40^, a voxel-wise non-parametric permutation t-test, using family-wise error correction and TFCE (threshold-free cluster enhancement) to consider for multiple comparisons. As a result, we obtained the significance map, containing voxel-wise corrected p-values. A threshold of corrected p<0.01 was applied to identify clusters of significant differences.

## ACKNOWLEDGMENTS

We are very grateful to Maria Teresa Muñoz, Ana Maria Lopez, Silvia Artigas and Albert Coll for excellent technical support. We are also grateful to the staff of the Confocal Microscopy Service and the Animal Experimental Unit of the Scientific and Technological Centers of the University of Barcelona (CCiTUB). We are indebted to the Magnetic Resonance Imaging Core Facility of the Institut d’Investigacions Biomèdiques August Pi i Sunyer for the scientific technical support in MRI acquisition and analysis.

## AUTHOR CONTRIBUTIONS

S.C-B, M.J.R., J.A., and M.M., contributed to the conception and design of the study and designed the experiments. S.C-B, L.S-R, E.G-G, L.G-G, A.S-B, S.F-G, E. R-U, A.G. J L-G, E.M-M, A.C and G.S performed all the experiments. S.C-B, E.M-M, A.C, G.S and M.M. performed the statistical analysis. J.A., M.M., A.G. and M.J.R. obtained financial support. S.C-B and M.M wrote the manuscript. All authors have reviewed and corrected the manuscript; they also agreed to the published version of the manuscript.

## DATA AVAILABILITY STATEMENT

The datasets used and/or analyzed during the current study are available from the corresponding author upon reasonable request.

## FIGURE LEGENDS

**Figure S1. Validation of D1-Cre and Adora2a-Cre mice and selection of optogenetic stimulation protocol.** (A) Heterozygous Drd1a-Cre+/- (D1) and Adora2a-Cre+/- (A2a) male mice were crossed with homozygous Flox-TdTomato female to validate pathway specificity, resulting in 50% D1-TdTomato and 50% A2a-TdTomato mice. Tdtomato was constitutively expressed in all D1- and A2a-expressing cells (cyan). Within the striatum (STR), D1-expressing cells (left) project to the substantia nigra pars reticulata (SNr) and give rise to the direct pathway, whereas A2a-expressing cells (right) project to the globus pallidus externus (GPe) and give rise to the indirect pathway. (B-C) Optogenetic stimulation was delivered unilaterally to the DLS, and induction of circling behavior analyzed by the number of contralateral and ipsilateral turns/ min induced, using a 473 nm laser at (B) constant light stimulation, as in Kravitz et al 2010 and at (C) 10 Hz light stimulation, as in Fernandez-Garcia et al 2020. Data are represented as mean ± SEM. Number of mice per group: D1-Cre n = 4 and A2a-Cre n = 2 mice. *p<0.05, ***p<0.001.

